# A comparative study of techniques for differential expression analysis on RNA-Seq data

**DOI:** 10.1101/005611

**Authors:** Zong Hong Zhang, Dhanisha J. Jhaveri, Vikki M. Marshall, Denis C. Bauer, Janette Edson, Ramesh K. Narayanan, Gregory J. Robinson, Andreas E. Lundberg, Perry F. Bartlett, Naomi R. Wray, Qiong-Yi Zhao

## Abstract

Recent advances in next-generation sequencing technology allow high-throughput cDNA sequencing (RNA-Seq) to be widely applied in transcriptomic studies, in particular for detecting differentially expressed genes between groups. Many software packages have been developed for the identification of differentially expressed genes (DEGs) between treatment groups based on RNA-Seq data. However, there is a lack of consensus on how to approach an optimal study design and choice of suitable software for the analysis. In this comparative study we evaluate the performance of three of the most frequently used software tools: Cufflinks-Cuffdiff2, DESeq and edgeR. A number of important parameters of RNA-Seq technology were taken into consideration, including the number of replicates, sequencing depth, and balanced *vs.* unbalanced sequencing depth within and between groups. We benchmarked results relative to sets of DEGs identified through either quantitative RT-PCR or microarray. We observed that edgeR performs slightly better than DESeq and Cuffdiff2 in terms of the ability to uncover true positives. Overall, DESeq or taking the intersection of DEGs from two or more tools is recommended if the number of false positives is a major concern in the study. In other circumstances, edgeR is slightly preferable for differential expression analysis at the expense of potentially introducing more false positives.

## Introduction

High-throughput cDNA sequencing (RNA-Seq) has emerged as an attractive and cost-effective approach for transcriptome profiling due to ongoing increases in throughput and decreases in costs of next-generation sequencing (NGS). Compared to microarray techniques, RNA-Seq can be performed without prior knowledge of reference sequences and enables a wide range of applications including transcriptome *de novo* assembly [1-4], abundance estimation [5-9], and detection of alternative splicing events [10-12], all of which have revolutionized our understanding of the extent and complexity of eukaryotic transcriptomes [13].

In RNA-Seq experiments, the primary interest of biologists in many studies is differential expression analysis between groups. To quantify gene expression, RNA-Seq reads need to be aligned to the reference genome for model organisms (e.g. human, mouse) or to the transcriptome sequences reconstructed using *de novo* assembly strategies for organisms without reference sequences. The number of mapped reads is calculated based on the outcome of the alignment to estimate the relative expression level of genes and subsequently statistical methods are applied to test the significance of differences between groups. The general workflow for the analysis of differential expression is illustrated in Figure 1.

**Figure 1.**
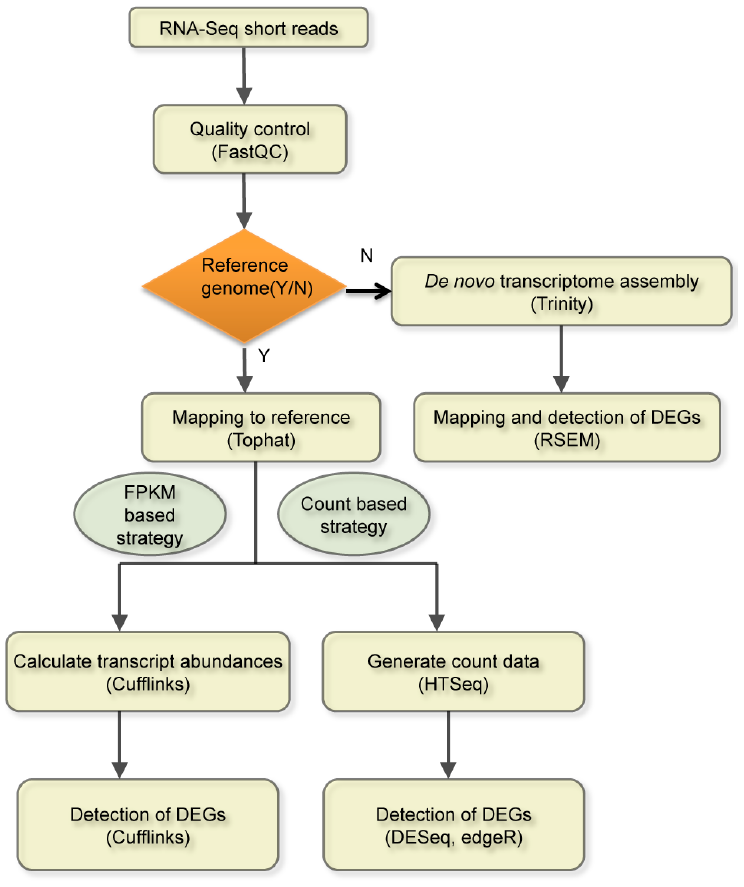
The workflow of differential expression analysis for RNA-Seq data.

Although initially it was claimed that RNA-Seq applications could produce unbiased, ready-to-analyze gene expression data [14,15], in reality it is nontrivial to accurately quantify gene expression and detect differentially expressed genes (DEGs). Difficulties faced by researchers in RNA-Seq study design and analysis are 1) general biases and errors inherent in the NGS technology (e.g. biases introduced during library preparation, nucleotide-specific and read-position specific biases in sequence quality and error rate) [16,17]; 2) biases of abundance measures due to the effects of nucleotide composition and the varying length of genes or transcripts [5]; 3) undetermined effects of both sequencing depth and the number of replicates; 4) the combination of technical and biological variation as well as biases within and between treatment groups that make it difficult to accurately discriminate real biological differences between groups; 5) the existence of alternative gene isoforms and overlapping sense-antisense transcripts may also compound difficulties in differential expression analysis [18].

Several attempts have been made to address the aforementioned difficulties (e.g. ref.16-18). In early RNA-Seq studies lacking biological replicates, the distribution of feature counts across technical replicates was reported to fit well to a Poisson distribution where the variance is equal to the mean [15,19]. However, when biological replicates are included, it has been noted that the Poisson distribution underestimates the variation seen in the data [20,21], a problem known as overdispersion. Therefore, the resulting statistical test based on Poisson assumptions does not control type-I error as advertised. As economical considerations typically do not allow large numbers of biological replicates, the Negative Binomial (NB) distribution has been proposed because of its ability to deal with the overdispersion problem [6,22]. With the introduction of a scaling factor for the variance, NB outperforms Poisson and currently it has achieved a dominant position in the methodologies to model feature counts for RNA-Seq data [5,6,22].

A number of tools have been developed that use the NB distribution for differential expression analysis based on RNA-Seq data (e.g. Cufflinks [8,18,23], DESeq [6], edgeR [22], etc.). Taking advantage of these tools, the power of the RNA-Seq approach to detect DEGs has been recently demonstrated [24-27]. However, there is no consensus on which analysis method is optimal, nor on how to approach a proper analysis to ensure validity of outcomes in terms of reproducibility, accuracy and robustness. Moreover, there is no agreement on how to design an efficient RNA-Seq study to maximize reward by balancing the amount of sequencing reads for each sample and the number of replicates for each group given a fixed budget.

Recently, several published studies have evaluated the performance of RNA-Seq software tools for differential expression analysis, although they all confined their analyses to methods that are applicable to count matrices (also known as count tables that contain raw counts for each feature) [28-30]. One of the most popular RNA-Seq tools, Cufflinks, does not use count matrices and has not been included in these comparisons. The dearth of comparisons may reflect that Cufflinks takes BAM files (the binary version of sequence alignment data) as input, which may make comparisons involving Cufflinks-Cuffdiff2 more cumbersome due to their large file sizes and binary format. In contrast, other methods such as DESeq and edgeR use count matrices as input, which are easier to handle and simulate for comparative evaluations.

Here, we evaluate the performance of Cufflinks-Cuffdiff2 as well as two widely used count matrices based tools - DESeq and edgeR, for DEG analysis of RNA-Seq data using both real RNA-Seq datasets and simulated datasets. In addition, we consider some important parameters of RNA-Seq technology that impact experimental design, including the number of biological replicates, sequencing depth, and balanced *vs*. unbalanced sequencing depth within and between groups. The results of this study will not only guide the choice of analysis tools, but more importantly will help researchers design optimal and efficient RNA-Seq experiments for their future studies.

## Material and Methods

### Ethics Statement

All animal treatments and experimental procedures were carried out in accordance with the Australian Code of Practice for the Care and Use of Animals for Scientific Purposes. The experimental procedures were approved by The University of Queensland Animal Ethics Committee (QBI/293/13/NHMRC).

### Tools for the differential expression analysis

The performance of Cufflinks-Cuffdiff2 [8,18], DESeq [6] and edgeR [22] are evaluated in this study. Cuffdiff2 is a program in the Cufflinks package (v2.1.1). It adopts an algorithm that controls cross-replicate variability and read-mapping ambiguity by using a model for fragment counts based on a beta negative binomial distribution. It can identify differentially expressed transcripts and genes, differential splicing and promoter-preference changes [18]. In this study we only focus on its function to identify differentially expressed genes (DEGs) so that we can compare its performance with that of the other two software packages. Both DESeq (version 1.10.1) and edgeR (version 3.0.8) are well-documented R/Bioconductor packages (release version 2.12 with R 2.15.1). DESeq implements a method based on the negative binomial distribution with variance and mean linked by local regression to detect DEGs [6]. EdgeR also models the variability of raw count data of RNA-Seq based on the negative binomial distribution. Empirical Bayes methods are used to moderate the degree of overdispersion across transcripts [22].

### RNA-Seq datasets

1. **The MAQC dataset:** MicroArray Quality Control Project (MAQC) data [31] includes biological samples for human brain reference RNA (hbr) and universal human reference RNA (uhr). Two different laboratories, Dr. Dudoit from University of California at Berkeley and Dr. Wu from Genentech, independently sequenced the samples using the Illumina GAII platform [32]. Dataset I from Dr Dudoit’s group contains 137.77 million (M) RNA-Seq reads with the read length of 35 bp whereas dataset II from Dr Wu’s group contains 112.70 M reads with the read length of 50 bp. Both datasets are single-end RNA-Seq reads without any biological replicates. In dataset I, hbr and uhr samples were sequenced using seven lanes across two flow cells, and detailed information of this dataset is given in Bullard et al.’s study [19]. This dataset was downloaded from NCBI sequence read archive (SRA) with ID SRX016359 and SRX016367. In dataset II, each sample was sequenced in seven lanes of one flow cell [32], the details regarding the dataset II are given in Nacu et al.’s study [33]. This dataset was downloaded from NCBI Gene Expression Omnibus (GEO) with ID GSE24284 (GSM597210 for hbr and GSM597211 for uhr). In both datasets, all reads were merged from different lanes for the same sample and the number of reads for each sample is shown in Table S1. The treatment groups compared for identification of DEGs are hbr *vs*. uhr.
2. **The K_N dataset:** The K_N RNA-Seq dataset was generated in-house at Queensland Brain Institute from mouse neurosphere cells treated by potassium chloride (KCl) and norepinephrine. Briefly, adult (8–12 week old) male C57BL/6J mice were used for all *in vitro* experiments conducted in this study. All mice were housed in groups and maintained on a 12-hour light/dark cycle with *ad libitum* access to food and water. After mice were sacrificed by cervical dislocation, the neural precursor activity was examined as described in detail in a previous study [34]. The isolated hippocampal tissue was minced to obtain a single cell suspension. The cell suspension was cultured in complete neurosphere medium containing EGF and bFGF, in the presence of norepinephrine (10 µM) or potassium chloride (15 mM). The primary neurospheres were collected on day 14 for each treatment. Total RNA was extracted from neurospheres using TRIzol Reagent (Life Technologies) and chloroform (Sigma-Aldrich) followed by precipitation and washing with isopropanol and ethanol respectively. RNA samples were treated with the Ambion DNA-free kit (Life Technologies) according to the manufacturer’s instructions to remove any contamination from genomic DNA. DNase-treated samples were assessed for their RNA integrity number (RIN) using an Agilent RNA 6000 Pico Kit (Agilent Technologies) on the Agilent 2100 Bioanalyser according to the manufacturer’s instructions. RNA samples with a RIN greater than 8 were selected and a total of 100 ng was used for each sample for sequencing. Total RNA was quantitated using the QuBit RNA assay kit (Life technologies, cat#Q32852). Sequence libraries were generated using the Illumina TruSeq RNA Sample Preparation Kit v2 (cat#RS-122-2001), with sample indexing/multiplexing. A total of four libraries (four biological replicates) were prepared for each of the treatment groups and sequenced using the Illumina Hiseq2000 platform. Data generated from potassium treatment group is referred to as “K”, and data from norepinephrine treatment is referred to as “N”, hence the “K_N” dataset. In total, 476.38 M paired-end reads with a read length of 101 bp × 2 were generated (231.55 M for K and 244.83 M for N), and we refer the full dataset of “K_N” as “K_N_full”. The data sizes of the four biological replicates in each group are shown in Table S2. In order to investigate the impact of sequencing depth on the performance of three software tools, we randomly subsampled four subsets of data with different sequencing depths from the “K_N_full” dataset. Each subset contains the balanced dataset around 30 M, 20 M, 10 M, 5M for each individual sample, and these subsets are named as “K_N_30M”, “K_N_20M”, “K_N_10M” and “K_N_5M”, respectively. In order to study the effects of unbalanced sequencing depth between treatment groups, another two subsets were generated. One subset includes 30 M of data in the “K” group but 5 M in the “N” group, and this subset is denoted as “unbalanced between groups 1”. The other unbalanced subset includes all data in the “K” group but 20 M in the “N” group and it is denoted as “unbalanced between groups 2”.
3. **The lymphoblastoid cell lines dataset:** The lymphoblastoid cell lines (LCL) RNA-Seq datasets were generated by Pickrell et al. [35]. It is a large population based RNA-Seq experiment, which sequenced 69 cell lines derived from Yoruban individuals from Nigeria by the International HapMap Project. Each sample was separately prepared and sequenced at two independent sequencing centers (the Yale sequencing center and the Argonne sequencing center) using the Illumina GAII platform.

In this study, we used samples with more than 8 M reads per individual from those sequenced at Yale to gain enough sequencing depth for each sample, which resulted in 40 samples in total (NCBI SRA accessions: SRR031811 - SRR031820, SRR031839, SRR031840, SRR031843 - SRR031846, SRR031848, SRR031857 - SRR031861, SRR031867 - SRR031875, SRR031877, SRR031893 - SRR031896, SRR031898, SRR031899, SRR031955, SRR031956). From these 40 individuals, we randomly selected and assigned N (N = 20, 14, 8, 6, 5, 4, 3, 2, 1) samples to each of the two hypothetical treatment groups. N is the number of biological replicates in each hypothetical treatment group. We expect no differentially expressed genes between the two hypothetical groups because samples were randomly selected from a same population. This dataset (hereafter referred to as the LCL1 dataset) is used to estimate false positive rates for the three software tools.

Comparisons of methods based on real RNA-Seq data are limited because a complete list of true DEGs and non-DEGs is not known. Therefore, we also used simulated data with known DEGs to estimate the true positive rate and false positive rate. Specifically, we simulated DEGs in two hypothetical groups using a strategy similar to that described in [29]. Briefly, DEGs were simulated to a random sample of 10% of the total genes. The count values of these DEGs were generated by scaling their original raw counts by exp{(−1)^i^δ_j_} where two different groups are indexed by i = 1, 2 and δ_j_ follows a two component normal distribution with parameter μ = (−0.5, 0.5) and σ = (0.7, 0.7). We employed this simulation design based on the LCL1 dataset. According to the simulated count values, bam files were generated by randomly adding or removing aligned reads using custom PERL scripts and SAMtools [36] for the application of Cuffdiff2. This dataset is named as the LCL2 dataset.

In addition, five randomly subsampled subsets (hereafter we referred as the LCL3 dataset) were generated based on the LCL2 dataset to test performance of each of three software tools using RNA-Seq datasets with different sequencing depths or with unbalanced sized data between/within groups, named as “S1_8M_balanced”, “S2_5M_balanced”, “S3_1M_balanced”, “S4_5M_1M_btw” and “S5_5M_1M_within”. Of them, “S1_8M_balanced”, “S2_5M_balanced” and “S3_1M_balanced” are subsets with around 8 M, 5 M and 1 M number of reads, respectively, for each sample in both conditions. “S4_5M_1M_btw” is the subset with numbers of reads being much higher in one condition (5 M) than that in the other condition (1 M). “S5_5M_1M_within” is the subset that the data size is unbalanced within each condition (i.e. numbers of reads for some of the samples in condition 1 and condition 2 are around 5 M while numbers of reads of other samples are around 1 M). See details in Table S3 for the information of each subset of LCL3 dataset. All studies for LCL1, LCL2 and LCL3 datasets were performed based on 10 independent simulations and the average values (i.e. number of DEGs, false positive rate and true positive rate) of the10 simulations were used.

### Benchmark datasets

1. **Quantitative RT-PCR data:** The results of quantitative RT-PCR (qRT-PCR) data from NCBI GEO (series ID GSE5350) was used as gold standard to evaluate the performance of three tools based on MAQC RNA-Seq datasets. We downloaded the four technical replicates of the uhr sample from GSM129638 to GSM129641, and the four technical replicates of the brain sample from GSM129642 to GSM129645, with a total of 1044 genes in the list. Among these genes, only those genes with the unique matched Refseq (hg19) gene ID were kept and genes with zero read counts in all samples of qRT-PCR were filtered out. The log2 ratio of fold change (log_2_FC) of the gene expression value between hbr and uhr was calculated according to Bullard et al.’s study [19]. Genes with |log_2_FC| > 2 (more than 4 fold differentially expressed between hbr and uhr) are considered as differentially expressed (the positive set). In contrast, genes with |log_2_FC| < 0.2 are considered as the negative set. Among the 1044 genes, 410 genes are in the positive set and 86 genes are in the negative set.
2. **Microarray data:** Microarray data for the neurosphere samples was generated at Queensland Brain Institute. Briefly, RNA was isolated from neurospheres generated in presence of either N or K (n = 3 each, two biological replicates in common with the RNA-Seq data) and converted to cDNA using Applause WT-Amp ST kit (NuGEN). Fragmentation and biotin labelling of the cDNA was carried out using Encore Biotin module (NuGEN) for Affymetrix GeneChip Mouse Gene 1.0 ST arrays. Labelled cDNA was hybridised to arrays in an oven at 45°C. Hybridised arrays were scanned using the Affymetrix GeneChip Scanner and the scanned data (.CEL files) was transferred to the Partek Genomics Suite and data analysis was carried out as described in Narayanan et al.’s study [37]. The results of cDNA microarrays have been employed to verify the performance of three software programs based on K_N RNA-Seq datasets. Among total 35557 genes of K_N microarray data, only genes with the unique matched Refseq (mm10) gene ID were retained. Genes with |log_2_FC| >1 and the corresponding *P*-value less than 0.05 were chosen as the positive set whereas genes with the |log_2_FC| < 0.1 and the *P*-value larger than 0.1 were chosen as the negative set. In total, we obtained 77 genes in the positive set and 9072 genes in the negative set from this microarray benchmark data.

### Differential expression analysis for RNA-Seq datasets

Our basic analyses followed the general workflow depicted in Figure 1. Briefly, Tophat (version 2.0.8) [38] was used to align the short reads to the reference human genome (hg19) for MAQC and LCL datasets and mouse genome (mm10) for K_N dataset. Cuffdiff2 was then used to generate the list of DEGs. For input into the DESeq and edgeR packages, the raw count table was produced by HTseq-count (version 0.5.4p2) and then the lists of DEGs were generated as recommended in the manuals of these two packages. An example of detailed commands for the differential expression analysis is listed in File S1.

### Evaluation of the performance of three techniques for detecting DEGs

The area under the receiver operating characteristic (ROC) curve (AUC) is used to evaluate the performance of different techniques for the detection of DEGs as suggested in [32]. For the MAQC RNA-Seq data, we compared the results of each software program with the positive and negative sets from qRT-PCR. For the K_N dataset, we compared the results from the full dataset and its subsets with the positive and negative sets from the microarray data. For the LCL dataset, since we introduced DE genes by simulation, a full set of known DEGs and non-DEGs could be used to evaluate the performance of three methods. The area under the receiver operating characteristic (ROC) curve (AUC) is used to evaluate the performance of different techniques for the detection of DEGs as suggested in [32]. Thus, true positives (TP), false positives (FP), true negatives (TN) and false negatives (FN) are calculated based on the comparison results between RNA-Seq datasets and the positive and negative sets of the corresponding benchmark dataset. The false positive rate (FPR) and the true positive rate (TPR) are defined as follows: FPR = FP/(FP + TN), TPR = TP/(TP + FN), (see also Table S4).

The relationship between TPR and FPR is depicted by the ROC. As pointed out by [32], two kinds of AUC can be calculated in our experiments: (1) AUC1 is the area under the ROC curve in the full range of FPR, i.e. 0 ≤ FPR ≤ 1 and the maximum AUC1 is 1; (2) AUC2 is the area under the ROC curve in the range of 0 ≤ FPR ≤ 0.05 and the maximum AUC2 is 0.05. AUC1 reflects overall performance of identification of DEGs for each tool. In contrast, AUC2 reflects the relative performance when we limit the range of the false positive rate to that often applied in discovery analyses.

### Data and source codes availability

The K_N RNA-Seq dataset has been deposited in the NCBI SRA with accession ID SRX516577. The microarray data for the neurosphere samples has been deposited in the NCBI Gene Expression Omnibus with accession ID GSE57440.

Analysis pipelines with detailed descriptions for subsampling datasets and generating BAM files based on simulated count values are listed in File S1. All custom PERL scripts mentioned in our analysis pipelines can be downloaded from GitHub https://github.com/Qiongyi/RNA-Seq-comparison/.

## Results and Discussion

### Biological and technical replicates

We first investigated the impact of biological and technical replicates on the performance of differential expression analysis for each of three tools. The MAQC RNA-Seq datasets include two technical replicates that were independently generated in two different laboratories. The K_N RNA-Seq dataset as well as its subsets include 4 biological replicates. The LCL RNA-Seq dataset as well as its subsets include 20 biological replicates in each group. All datasets have been processed through the analysis pipeline shown in Figure 1 (reference genome available).

As expected, analyses from all three software tools with biological replicates or technical replicates outperform those without replicates as judged by both AUC1 and AUC2 (Figure 2 A-I). Interestingly, the performance of edgeR seems to be independent of the number of replicates for the MAQC dataset (Figure 2 C) but not for the other two datasets (Figure 2 F and I), which can be explained as follows. A parameter, the biological coefficient of variation (BCV), has to be set manually when there are no replicates. As documented [8], the BCV for datasets arising from well-controlled experiments is about 0.4 for humans. We thus set the BCV value to 0.4 when we employed edgeR to analyze the MAQC dataset and it seems that 0.4 is ideal for the BCV value for this MAQC dataset. On the other hand, when edgeR was employed in the context of no biological replicates for the K_N dataset, we set BCV to be 0.4, but for this mouse dataset the performance of edgeR is poor (Figure 2 F, blue solid line). After optimizing the BCV to 0.2, the performance was dramatically improved (Figure 2 F light blue dotted line), though not perfectly matching the performance of conditions with replicates. It demonstrates that achieving the optimum BCV value is critical for the performance of edgeR when there are no replicates available.

**Figure 2.**
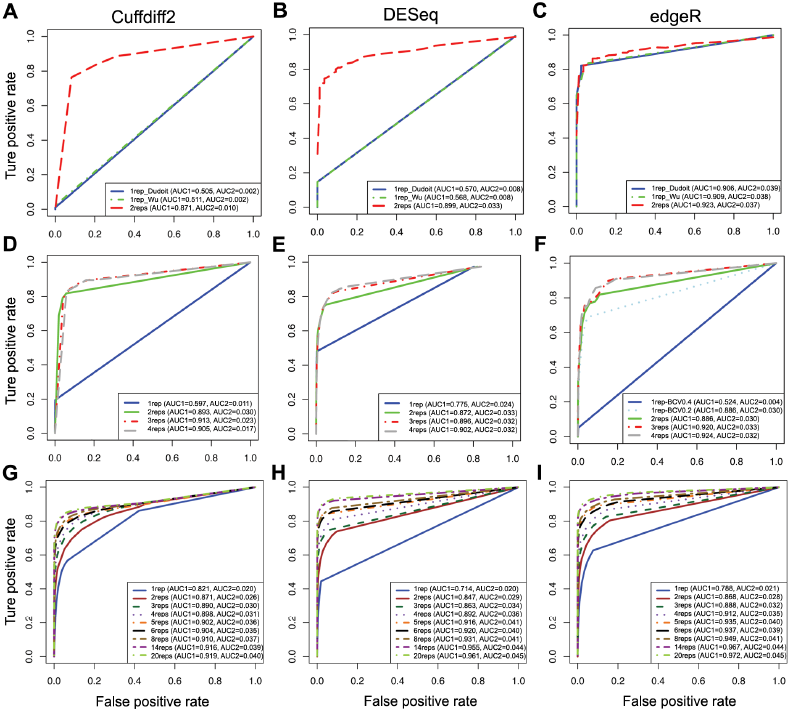
The effects of replicates for detecting DEGs based on ROC curves. ROC curves for evaluating the performance of Cuffdiff2, DESeq and edgeR on 1 to 2 technical replicates based on the MAQC dataset (A-C), 1 to 4 biological replicates based on the K_N dataset (D-F), and 1 to 20 biological replicates based on the LCL2 dataset (H-I).

For the K_N dataset, the incremental improvement became smaller as judged by the AUC as the number of biological replicates increased (Figure 2 D-F). However, for the LCL dataset, we always see an obvious improvement with increasing number of replicates until N = 14 replicates (Figure 2 G-I). With an increasing number of biological replicates, we expect more statistical power for the identification of DEGs, and a positive correlation between DEGs and the number of biological replicates is found for all three tools (Figure 3 A-B). Indeed, the correlation is almost linear between the number of detected DEGs and the number of biological replicates for DESeq and edgeR using the K_N dataset, whereas a different pattern is observed when using the LCL2 dataset, which indicates that the power for the detection of DEGs reflects both the amount of biological variation in the samples and the number of biological replicates. Based on the LCL2 dataset, the number of DEGs detected by Cuffdiff2 increases steadily with the number of replicates but with an obvious drop in the number of detected DEGs with 5 biological replicates (Figure 3 B). In contrast, a previous study reported that, with Cuffdiff2, the number of detected DEGs decreased when the number of samples increased [39]. These differences may be due to the fact that different versions of Cuffdiff2 were used between the studies and this could affect the performance of Cuffdiff2, which is also mentioned in Seyednasrollah et al.’s study [39].

**Figure 3.**
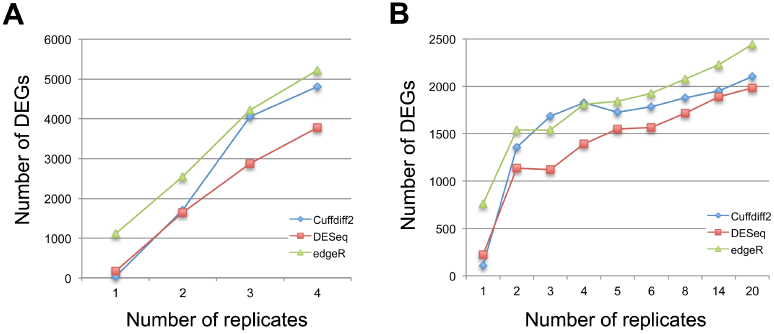
The effects of biological replicates on differential expression analysis. The numbers of differentially expressed genes identified by each of three tools under different numbers of biological replicates based on the K_N dataset (A) and the LCL2 dataset (B).

Generally, although each tool adopts different underlying statistical models to identify DEGs from RNA-Seq data, they all need replicates to separate the real differences between treatment groups from the random variation within groups. In the LCL dataset, when the amount of biological variation in the samples is large, all three tools take advantage of biological replicates to gain better performance. Recent studies [39,40] also demonstrated the importance of biological replicates for current RNA-Seq analysis tools to detect real biological differences in the presence of random biological variation, with Seyednasrollah et al. reporting that the SAMseq method exhibited most sensitivity to the number of replicates [39].

### The effects of the sequencing depth

To determine how sequencing depth affects the performance of differential expression analysis for the three software tools, we randomly subsampled seven subsets of data with different sequencing depths from the K_N and LCL datasets. The results of experiments from both the K_N and LCL datasets show that Cuffdiff2 is the most sensitive to sequencing depth while the performance of DESeq and edgeR are stable under different sequencing depths (Figure 4 A-F). It suggests that DESeq and edgeR are a better choice than Cuffdiff2 for differential expression analysis when sequencing depth is low (i.e. number of reads < 10 M). The results here also indicate that 20 M reads are sufficient for DEGs analysis using Cuffdiff2, which is a relatively cost-effective sequencing depth. However, as expected, the number of DEGs discovered by each tool decreased with a reduction in sequencing depth (Figure 5 A-B), which is in line with a previous study [41]. EdgeR identifies more DEGs (Figure 5) but also has a higher FPR in simulated data (see below).

**Figure 4.**
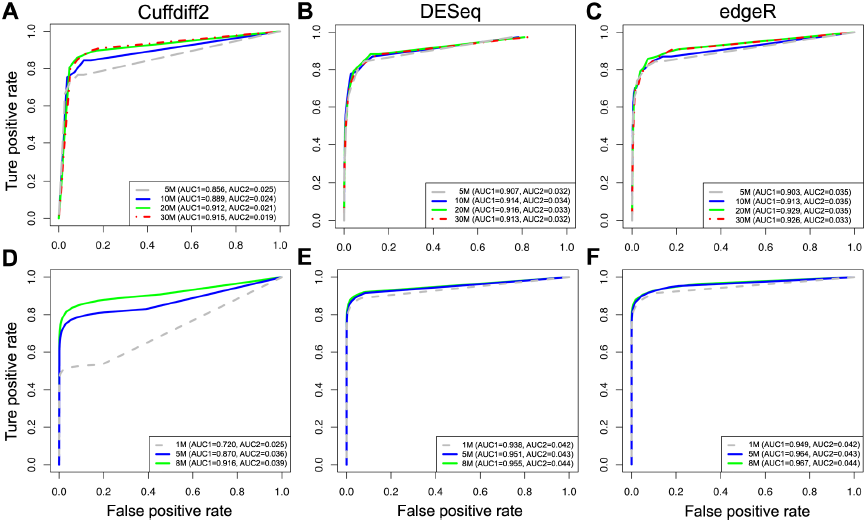
The effects of sequencing depth for detecting DEGs. ROC curves for evaluating the performance of Cuffdiff2, DESeq and edgeR with different sequencing depths based on the K_N subsets (A-C) and the LCL3 simulated dataset (D-F).

**Figure 5.**
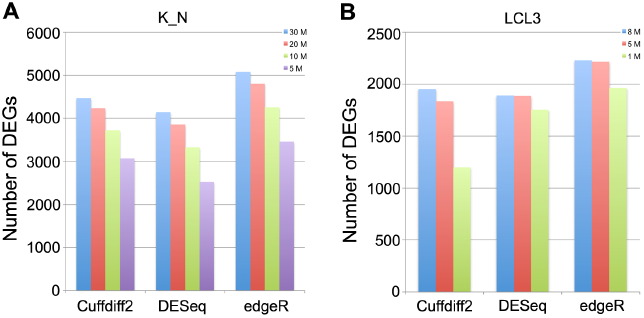
The effects of sequencing depth on differential expression analysis. The numbers of differentially expressed genes identified by Cuffdiff2, DESeq and edgeR are shown based on the K_N subsets (A) and the LCL3 simulated dataset (B).

### The effects of unbalanced sequencing depth

In order to understand the impact of unbalanced sequencing depth on the performance of each tool for differential expression analysis, we made comparisons based on subsets of data from the K_N and LCL datasets with balanced sequencing depth among all samples, unbalanced sequencing depth within groups and unbalanced sequencing depth between groups. The sequencing depth of the K_N_full dataset is unbalanced within groups, as shown in Table S2, with the data sizes ranged from 30 M to 116 M.

We observed from both K_N and LCL datasets that DESeq is (marginally) most sensitive to unbalanced sequencing depth between groups (Figure 6). AUC1 of DESeq decreases and AUC2 slightly decreases when using unbalanced subsets (Figure 6 B and E). Cuffdiff2 is more sensitive to overall sequencing depth rather than unbalanced sequencing depth between groups (Figure 6 A and D). EdgeR has the maximum AUC1 and AUC2 under these scenarios without obvious negative effects from unbalanced sequencing depth (Figure 6 C and F).

### Trade-off between sequencing depth and number of replicates

Since budgetary constraints are common with sequencing, optimal experimental designs need to balance the sequencing depth for each sample with the number of replicates for each group. As our results show (Figures 2 and 4) and as others have reported [40,42], the overall impact of the sequencing depth is not as critical as the number of biological replicates. Including sufficient biological replicates should be a prime consideration for RNA-Seq study designs, and determining a suitable number of biological replicates is a key question. Based on overall performance as judged by AUC1, in the controlled groups of our mouse neurosphere project our results suggest that N = 3 or 4 replicates is sufficient (Figure 2 D-F), but the number of detected DEGs continues to increase linearly as the number of replicates increases (Figure 3 A), although our expectation is that the linear increase will not continue. For example, Seyednasrollah et al. showed that the number of DEGs plateaus at 4-6 replicates (samples from inbred mice) using tools such as DESeq, edgeR, baySeq, limma, etc. [39]; their study may be similar to our K_N dataset. In contrast, we find in the LCL data derived from unrelated Yoruban individuals that although the number of detected DEGs changes little after N = 4 replicates (Figure 3 B), the overall performance continues to increase (as judged by AUC1 in all three tools, Figure 2 G-I) as the number of biological replicates increases, presumably reflecting increasing accuracy of the detection of DEGs. This indicates the importance of biological replicates to uncover real biological differences when random biological variation within groups is relatively large. Under a fixed budget, the cost of generating biological replicates must be factored into the experimental design, and on balance, we advise maximizing of the number of biological replicates within these constraints.

**Figure 6.**
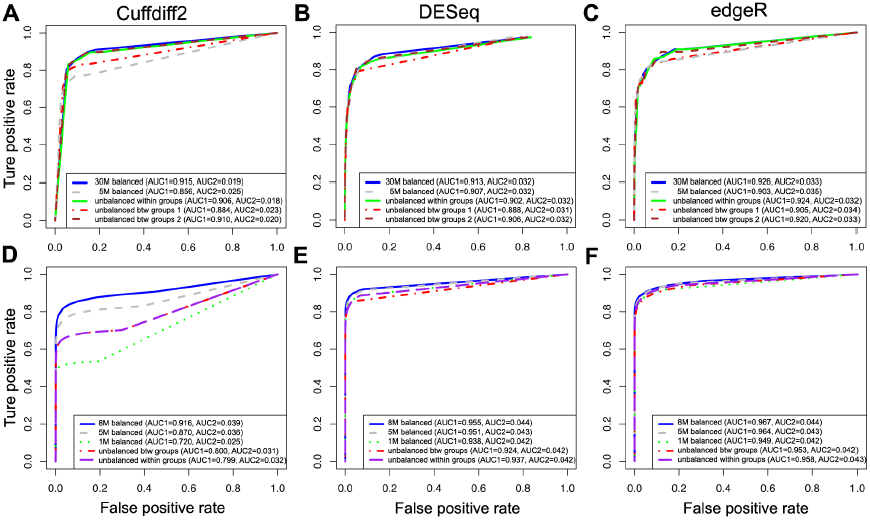
The effects of balanced and unbalanced depths of reads for detecting DEGs based on ROC curve. ROC curves for evaluating the performance of Cuffdiff2, DESeq and edgeR on balanced and on unbalanced depths of reads based on the K_N dataset (A-C) and the LCL3 simulated dataset (D-F).

Our results suggest that more biological replicates are required to identify DEGs using RNA-Seq when studying human cell lines from unrelated individuals in the same ethnic group than equivalent studies based on neurosphere cells from inbred mice. This suggests that the amount of biological variation in the samples to be sequenced should be a major consideration when determining the optimal number of biological replicates required for a given RNA-Seq study. It is likely that more biological replicates will be required in animal/human tissue samples compared to human cell lines or cells from inbred lab strain models. However, more gold standard datasets for benchmarking and more comprehensive evaluations based on these datasets are required to guide future RNA-Seq study designs in terms of the optimal number of biological replicates relative to the amount of biological variation in the samples.

### The performance of three tools

For all three datasets, we find that edgeR performs slightly better than the other two tools as judged by the two AUC statistics (Figure 7 A-C). A recent update of the Cufflinks software package (version 2.1) introduced a new testing method into the Cuffdiff2 program that aimed to substantially improve performance over previous releases. For comparison we also evaluated the pre-update version of the Cuffdiff2 program (v2.0.2). We find that the versions do perform differently under our test scenarios both in terms of the AUC statistics (Figure 7 A-C) and the number of identified DEGs (Figure S1). Surprisingly, we noticed that the performance of Cuffdiff2 (v2.0.2) was inversely correlated with the number of biological replicates in our LCL dataset when the number increased from 3 to 14, and the best performance as judged by AUC1 and AUC2 was achieved when there are 3 to 5 biological replicates (Figure S2). The overall performance of Cuffdiff2 for differential expression analysis at gene level resolution is not as good as that of the two count based tools evaluated in this study, as judged by both AUC1 and AUC2 (Figure 4 and Figure 7 A-C). One qualification is that Cuffdiff2 achieves similar performance compared with DESeq when there is sufficient sequencing depth (i.e. ≥ 20 million reads) for each individual sample (Figure 4 A and B). A possible explanation is that Cuffdiff2 transforms the alignment results to FPKM (Fragments Per Kilobase of gene model per Million fragments mapped) values rather than to raw count values. It then performs statistical tests based on the beta negative binomial model, under the assumption that this reflects the underlying distribution of the FPKM. However, FPKM, or its counterpart RPKM (Reads Per Kilobase of gene model per Million reads mapped) when a single-end sequencing strategy is applied, may not be an appropriate way to normalize RNA-Seq data, as discussed in many studies [43,44]. For example, FPKM normalization has been shown to reduce sample variability when compared with raw counts, with the resulting bell-shaped curve slightly skewed towards negative values rather than centered over a zero value [44]. A more recent study also indicated that transcript length normalization may not be a good strategy and may cause conservative bias in Cuffdiff2 [45].

**Figure 7.**
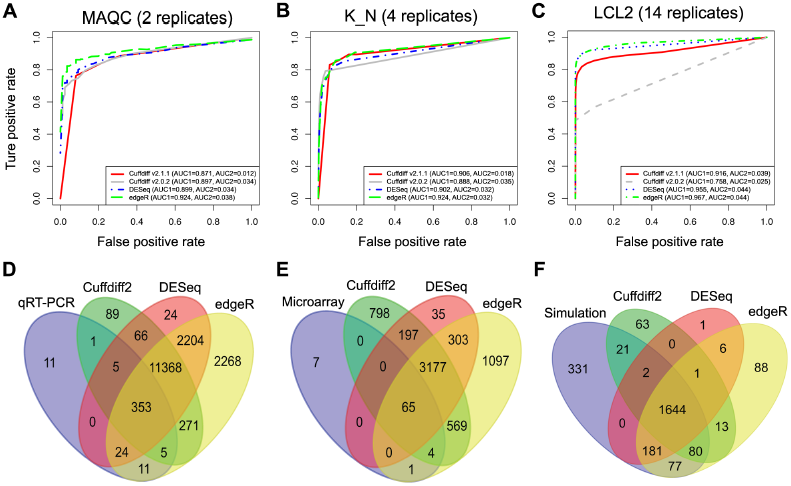
The performance of the three tools. ROC curves of the three tools are shown based on the MAQC (A), K_N (B) and LCL2 (C) datasets. Venn diagrams are used to show the intersection of the numbers of the differentially expressed genes identified by three tools compared with the benchmarks based on MAQC (D), K_N (E) and LCL2 (F).

In this study, although most of the DEGs identified by each of three tools overlapped, edgeR always detected more DEGs than the other tools, with a number of DEGs identified exclusively by edgeR (Figure 7 D-E). However, identification of more DEGs is not always a good thing, since it may introduce more false positives that make the experimental validation difficult. As an example shown in the comparison between DEGs identified by edgeR and true DEGs generated by simulation, edgeR introduced the most (108) false positives compared with Cuffdiff2 (77) and DESeq (8) (Figure 7 F). Using the LCL1 dataset, where we expect no differentially expressed genes between groups, we found all three tools performed well with reasonably small false positive rates. DESeq achieved the best performance in this scenario with the least number of false positives when there were biological replicates available (Table S5).

It is interesting to note that, although both DESeq and edgeR are negative binomial distribution model-based algorithms using the same count matrices as input, many genes with high fold change were picked up by edgeR but not by DESeq (Figure S3). When we looked into these particular DEGs, we observed that these DEGs were either lowly expressed (small raw read count values) or highly variable amongst biological replicates (i.e. high variance within groups) (File S2), indicating that DESeq is more conservative than edgeR in identifying DEGs under these conditions. This is in line with findings from previous studies [28,29] which found that DESeq is more conservative than other evaluated count based tools. We also observed that DEGs exclusively identified by either DESeq or edgeR showed a broader range of fold change than that of Cuffdiff2-exclusive DEGs (Figure S3). Since Cuffdiff2 is optimally designed to detect differential expression at the transcript level based on different underlying models and assumptions compared to DESeq and edgeR, it is not surprising that there are differences in the statistical significance testing for the identification of DEGs.

### Limitations

Our study has several limitations that may constrain generalization of results. Firstly, we considered only three experimental designs, one with two technical replicates, one with four biological replicates, and one from simulation based on real RNA-Seq data with 20 biological replicates. Comparisons may need to be made over a broader range of scenarios to draw general conclusions. Secondly, to determine true and false positive rates of the tools we had to benchmark against a “gold standard” for real RNA-Seq data. Here the DEGs from the benchmarked data sets were obtained from qRT-PCR or microarray analyses. However, false positives may exist in these lists. For the mouse neurosphere project only 77 genes were selected as a positive set from the microarray results, which may mislead the interpretation when compared with RNA-Seq results. Considering this, we attempted to overcome these limitations by also using simulated data sets in which DEGs were known. Thirdly, in the mouse neurosphere project two of the three biological replicates in the microarray experiment were in common with the RNA-Seq analysis. A cleaner interpretation would be achieved by the benchmarked set being either fully the same or fully different. Fourthly, there are several other count matrices based tools available for the differential expression analysis, such as NBPSeq [46], baySeq [47], SAMSeq [48] and ShrinkSeq [49], among others, but a thorough comparison of all these tools is beyond the scope of the present study. Despite these limitations, data sets that allow benchmarking in this way are relatively rare and so we draw the best conclusions within these constraints.

## Conclusions

In this study, we conducted a comprehensive investigation to evaluate the performance of three of the most widely used software tools (Cuffdiff2, DESeq and edgeR) for differential expression analysis of RNA-Seq data while considering a number of important parameters of RNA-Seq experiments, including number of replicates, sequencing depth, and the unbalanced data within or between groups. By using the results from qRT-PCR and microarrays as benchmarks, we observed that edgeR performs better than DESeq and Cuffdiff2 in terms of the ability to uncover true positives with the default FDR setting (FDR < 0.05). However, this reflects that edgeR could always detect more DEGs than the other two tools (e.g. For the K_N dataset, 8% and 38% more than Cuffdiff2 and DESeq respectively), which may also introduce more false positives. All three tools perform much better when there are biological or technical replicates available. Consistent with previous studies it suggests that biological replicates are a key factor for differential expression analysis in RNA-Seq datasets [50]. The optimal number of biological replicates is strongly dependent on variability between biological replicates. In experiments designed with no replicates, edgeR is recommended but the value of BCV should be carefully set based on pilot data.

Our results show that Cuffdiff2 is most sensitive and DESeq is least sensitive to sequencing depth, but the overall impact of sequencing depth is not as critical as the number of biological replicates, which is in agreement with previous studies [40,42]. As our results indicate, the recommended sequencing depth for mouse RNA-Seq is around 20 M for each sample if Cuffdiff2 is to be used. When resources are limited, for the same number of total reads, an increased number of biological replicates each with reduced read depth is recommended over fewer replicates more deeply sequenced. In addition, DESeq is more sensitive to unbalanced sequencing depth between groups than the other methods. EdgeR has the best performance as judged by two AUC statistics without obvious negative effects under unbalanced sequencing depth.

No single method is clearly superior for differential expression analysis, since each has particular strengths that may be suitable for specific RNA-Seq datasets. Considering the overall performance based on three independent datasets in this study, Cuffdiff2 is not recommended for differential expression analysis at gene-level resolution, particularly if sequencing depth is low (i.e. < 10 million reads per individual sample). DESeq or the intersection of DEGs from two or more tools is recommended if the number of false positives is a major concern in the study, although DESeq should be avoided if sequencing depth is unbalanced between groups. Since titration of Illumina RNA-Seq libraries is difficult and multiplexing of samples is common (e.g. typically 3-12 libraries per HiSeq2000 lane), it is largely expected to have unbalanced library sizes or low sequencing depth for some samples. EdgeR can tolerate both of these factors and thus is slightly preferable for differential expression analysis at the expense of potentially introducing more false positives.

## Acknowledgements

The authors would like to thank QBI-IT for their assistance on computation support, Dr. Marie-Jo Brion and Dr. Jacob Gratten for discussions and critical reading of the manuscript, and the reviewers for their valuable comments and suggestions to improve the quality of the paper.

## Supporting Information

Table S1. Numbers of reads for the human hbr and uhr samples from the MAQC dataset. **(DOC)**

Table S2. Numbers of reads for the mouse neurosphere samples for treatment groups of K and N (the K_N dataset). **(DOC)**

Table S3. The number of reads for each individual sample of the LCL3 dataset. **(DOC)**

Table S4. The definition for TP, FP, TN, FN, TPR and FPR. **(DOC)**

Table S5. The false positive rate for Cuffdiff2, DESeq and edgeR based on the LCL1 dataset. (DOC)

Figure S1. Venn diagram showing the number of differentially expressed genes identified by two versions of Cuffdiff2. (TIFF)

Figure S2. The effects of biological replicates on the differential expression analysis for Cuffdiff v2.0.2. (TIFF)

Figure S3. The detected fold changes of all the differentially expressed genes identified by three tools were compared and shown, including DESeq *vs*. edgeR (top panel), DESeq *vs*. Cuffdiff2 (middle panel) and edgeR *vs*. Cuffdiff2 (bottom panel). (TIFF)

File S1. Analysis pipelines, methods and examples of commands for differential expression analysis, subsampling fastq files and generating SAM/BAM files based on simulated count values. (TXT)

File S2. The raw count values for genes with high fold changes were picked up by edgeR but not by DESeq. Genes with high fold changes (the absolute value of log2 fold changes larger than 2) identified as DEGs by edgeR but not by DESeq are listed in the file. The gene ID, the log2 fold changes (logFC) and FDR from DESeq, the logFC and FDR from edgeR, the raw count values for the four replicates of sample K (K1-K4) and sample N (N1-N4) are shown in each of the columns. (XLS)

